# Pseudohypoxia-stabilized HIF2⍺ transcriptionally inhibits MNRR1, a druggable target in MELAS

**DOI:** 10.1101/2024.10.07.617011

**Authors:** Neeraja Purandare, Vignesh Pasupathi, Yue Xi, Vikram Rajan, Caleb Vegh, Steven Firestine, Tamas Kozicz, Andrew M. Fribley, Lawrence I. Grossman, Siddhesh Aras

**Author notes:** To whom correspondence should be addressed: (L.I.G.); (S.A.); (A.M.F.).

## Abstract

The observation that amounts of the mitochondrial regulator MNRR1 (CHCHD2, AAG10, PARK22) are reduced in several pathologies, and that restoration of its level normalizes the pathological phenotype, prompted a search for compounds that could increase MNRR1 levels. High throughput screening of a 2400-compound drug and natural products library uncovered the antifungal drug nitazoxanide and its metabolite tizoxanide as effective enhancers of MNRR1 transcription. Using the mitochondrial disease MELAS (in which various mixtures, called heteroplasmy, of wild-type and mutant mitochondrial DNA (mtDNA) coexist) as a test, we showed that treating a cybrid MELAS model with tizoxanide could restore cellular respiration, enhance mitophagy, and, importantly, shift heteroplasmy toward more wild-type mtDNA. Furthermore, in MELAS patient fibroblasts, the compound could improve mitochondrial biogenesis, enhance autophagy, and protect the fibroblasts from LPS-induced inflammation. Chemical activation of MNRR1 is thus a potential strategy to improve mitochondrial deficits seen in MELAS. Investigation of the mechanism by which MNRR1 is reduced identified that two factors compete to regulate transcription at the MNRR1 promoter – RBPJκ, which stimulates it, and HIF2α, which inhibits it. In MELAS cells there is a pseudohypoxic state that stabilizes HIF2α, leading to transcriptional inhibition of MNRR1. Nitazoxanide reduces the levels of HIF2α by increasing the levels of PHD3, the prolyl hydroxylase that degrades HIF2α.

## Introduction

Mitochondria are well known to be genetic hybrids between the products of nuclear DNA (nDNA) and mitochondrial DNA (mtDNA) [1, 2]. Mutations in mtDNA, and its multicopy nature in which thousands of copies can be present in a diploid cell, give rise to a mixture of wild-type and mutant copies, termed heteroplasmy. Although there are diseases of homoplasmy, where only mutant mtDNA is present (*e.g.*, LHON [3]), many mtDNA diseases are heteroplasmic like MELAS (Mitochondrial Encephalomyopathy, Lactic Acidosis and Stroke-like episodes), a rare genetic disease that affects multiple organs. In such cases of a pathogenic mtDNA mutation, the higher the heteroplasmy level the more severe the phenotype [4, 5]. As a result, reducing the heteroplasmy level represents a viable treatment strategy. Doing so can involve increasing the amount of wild-type mtDNA, reducing the amount of mutant, or both.

Several pathogenic mtDNA mutations are specifically associated with the MELAS syndrome, the most common being m.3243A>G in the mitochondrial *MT-TL1* gene that codes for tRNA^Leu(UUR)^. The mutation causes hypophysiological mitochondrial protein translation and synthesis and can attain heteroplasmy levels >50% [6]. Mitochondria that cannot adequately translate and assemble electron transport chain (ETS) subunits produce insufficient energy to meet the requirements of their resident tissues along with other deficiencies, resulting in the characteristic MELAS multi-organ dysfunction whose sequelae are difficult to treat. For example, the overall energy deficiency stimulates mitochondrial proliferation in endothelial cells, precipitating the angiopathy and impaired blood perfusion that exacerbate organ damage and the stroke-like episodes. Deficiency of nitric oxide (NO), which regulates smooth muscle relaxation, contributes to these sequelae, including hypertension, fatigue, and memory loss. Treatments that increase NO levels are common clinical approaches currently under study. However, there are no specific standard treatments available to MELAS patients and many die between the ages of 10 and 35, underscoring a considerable unmet patient need.

Mitochondrial Nuclear Retrograde Regulator 1 (also called CHCHD2, PARK22, AAG10) is a biorganellar regulator of mitochondrial and nuclear function. MNRR1 was discovered in a computational screen to identify factors regulating the ∼90 proteins that comprise the oxidative phosphorylation complex [7]. Work since then has revealed that mitochondrial MNRR1 binds to cytochrome *c* oxidase (COX) to activate respiration [8, 9] and interacts with Bcl-xL to impede the extrinsic apoptosis cascade [10]. Nuclear located MNRR1, along with the protein RBPJκ, functions to activate transcription by binding to the conserved oxygen-responsive element (ORE) of its own promoter as well as to a host of other stress-response genes [11, 12]. Recent work from our group demonstrated that ectopic expression of MNRR1 could rescue the MELAS mitochondrial phenotype *in vitro* by increasing OXPHOS and the expression of CREBH target genes, which led to a significant decrease in heteroplasmy [12]. We also found that the transcription of MNRR1 was inhibited in MELAS cybrid cells. Thus, we sought small molecules that could activate transcription of *MNRR1* (and its target genes) to improve MELAS heteroplasmy and its associated OXPHOS pathologies. We identified nitazoxanide as an activator of MNRR1 transcription and uncovered a novel mechanism by which MNRR1 transcription is inhibited in MELAS and which is averted by the drug to enhance transcription.

## Results

### High throughput screen to identify MNRR1 activators

We have previously shown that activation of MNRR1 using exogenous overexpression rescues mitochondrial deficit in MELAS cybrid cells and stable overexpression was able to shift heteroplasmy towards wild-type (WT) mitochondrial DNA [12]. Hence, we were interested in identifying chemical activators of MNRR1 that could be repurposed as a therapeutic intervention in MELAS patients. To identify activators of MNRR1, we performed a screen of 2400 FDA-approved compounds using two independent cell lines, HEK293 and MDA-MB-468, stably expressing the *MNRR1*-promoter driven luciferase reporter (**Fig. 1A**). We selected compounds that activated the reporter by at least 50% or higher (**Fig. 1B**), identifying 54 and 155 compounds on the HEK293 and MDA-MB-468 screens, respectively (**Fig. S1A**). Thirteen compounds were common to both cell lines and, of these, 7 were selected based on their lower toxicity profile. From these, six were selected based on clinical availability (**Fig. 1C**). We then validated these compounds in a 143B osteosarcoma cell line (DW7) cybrid with ∼73% MELAS mutant mtDNA (m.3243A>G) in which MNRR1 levels are reduced [12] (**Fig. S1B**). We chose compound 4 since this was available as a clinical formulation that has been used for *in vivo* testing and shown to increase *MNRR1* transcripts and protein levels [13]. We also confirmed the increase of MNRR1 in one of the original cell lines used for identifying the activators – MDAMB468 (**Fig. 1D**) as well as in several other human cell lines (using tizoxanide, the active nitaxozanide metabolite – see below) (**Figures S1 C-E**).

**Figure 1:**
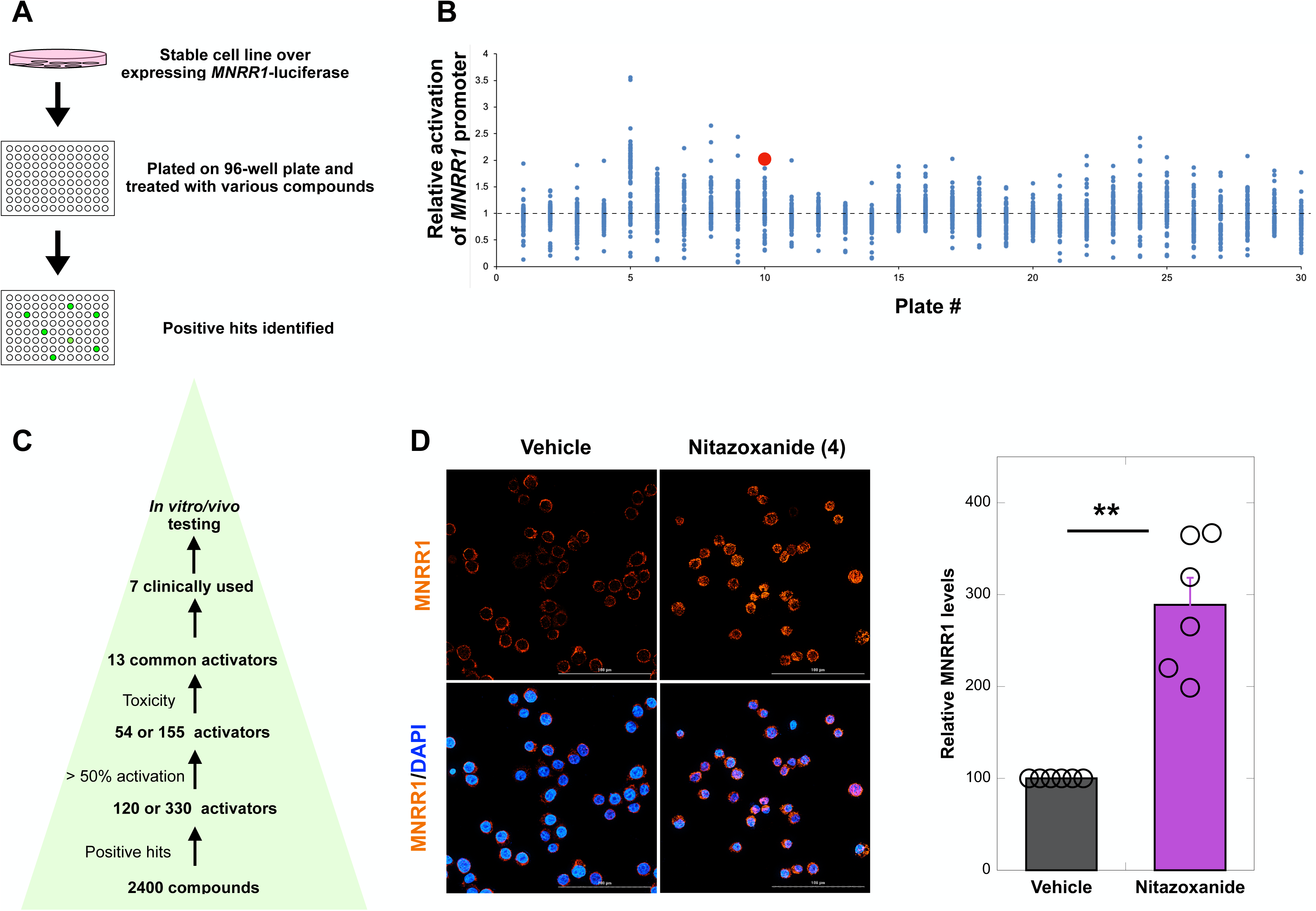
High throughput screen to identify MNRR1 activators. **A:** Overview of the high-throughput screening process to identify activators of MNRR1. **B:** Campaign view representing compounds that activate MNRR1. Control levels are represented as a dotted line. Each compounds is shown as a blue dot and nitazoxanide is highlighted as a red dot. **C:** Scheme for screening for strong activators of MNRR1 from high throughput screen. **D:** MDAMB468 cells treated with Vehicle (DMSO) or nitazoxanide (10 μM) for 24 h were immunostained (MNRR1=orange, DAPI=blue) and imaged at 63X using a confocal microscope. The scale bar represents 200 μm.

Nitazoxanide in cells is metabolized to tizoxanide (>97%) plus minor metabolites such as aminonitrothiazole and gentisate [14]. Only nitazoxanide-and tizoxanide-treated DW7 cells displayed a significant increase in the protein levels of MNRR1 (**Fig. S2A**). To confirm that the effects were at the transcriptional level, we measured *MNRR1* transcripts and observed that both compounds induced its transcription (**Fig. S2B)**.

### MNRR1 activation using tizoxanide enhances mitochondrial biogenesis and mitophagy to shift heteroplasmy in MELAS cybrid cells

We first confirmed the effects of MNRR1 activation by measuring oxygen consumption, which was increased (**Fig. 2A**). We had previously shown that MNRR1 overexpression induces homeostatic pathways such as mitophagy and mitochondrial biogenesis to aid in rescuing the phenotype. We had also shown that MNRR1 overexpressing cells display a reduction in heteroplasmy, making it an attractive therapeutic target for MELAS [12]. We therefore tested tizoxanide on MELAS cybrid cells and found it increased the proportion of WT mtDNA, by about 14% here, as shown by HaeIII digestion of the mtDNA fragment harboring the MELAS point mutation (**Fig. 2B**). Stable overexpression of MNRR1 in MELAS cybrid cells activated multiple homeostatic genes (**Fig. 2C**) and restored healthy mitochondria, presumably in part by stimulating mtDNA synthesis (**Fig. 2D**) by increasing PGC1⍺ (**Figs. 2C**, **E)** and enhancing mitophagy. Increased mitophagy is shown by increase of PINK1 (**Figs. 2C**, **E)**, autophagosomal proteins LC3A and LC3B (**Fig. 2E**), and by increased levels of the mitophagy marker pSer-65 ubiquitin (**Fig. 2F**) [15]. Taken together, these results suggest that tizoxanide increases MNRR1 levels and thereby induces the downstream pathways that rescue defective mitochondrial function in MELAS cybrid cells.

**Figure 2:**
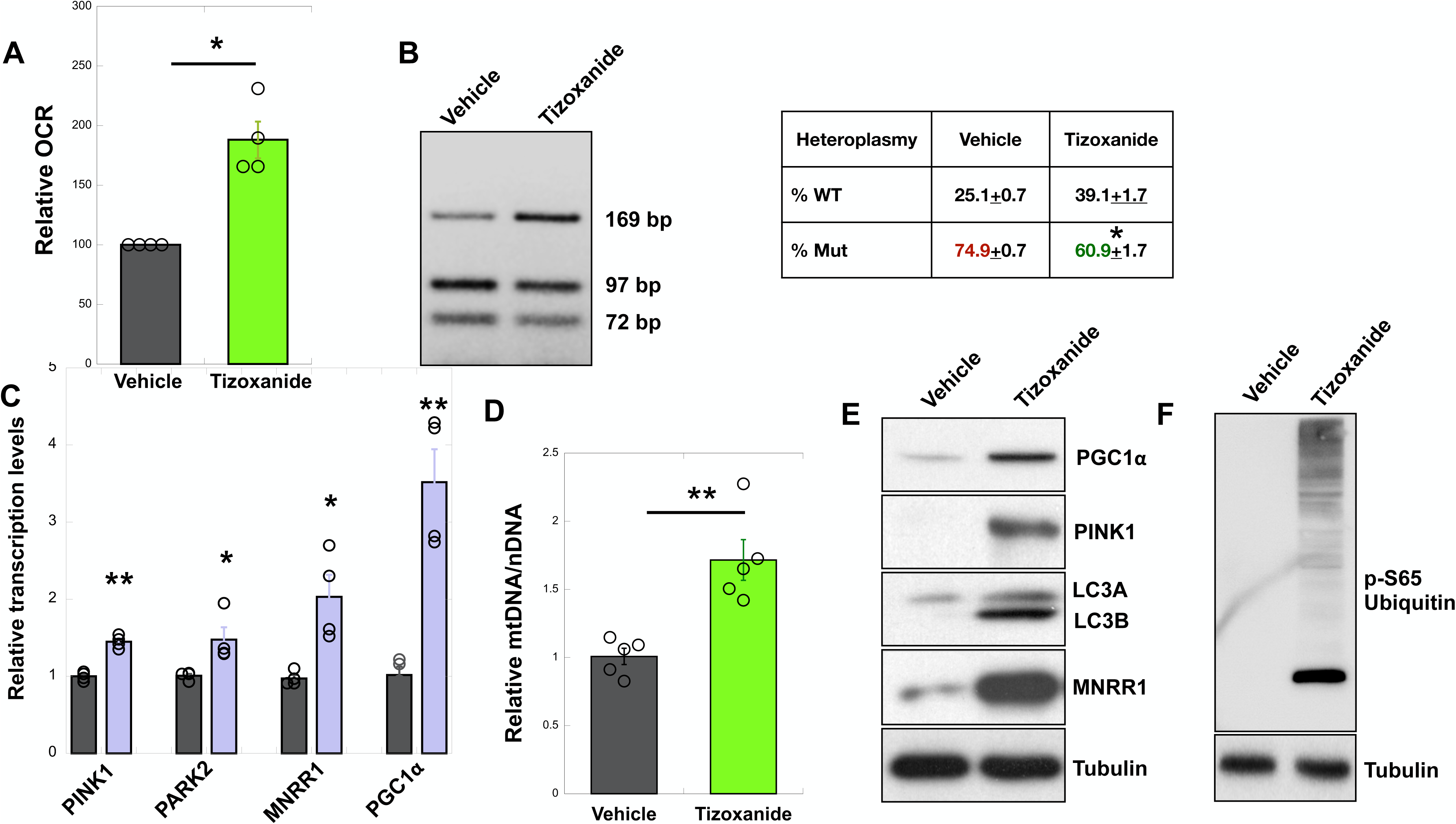
MNRR1 activation by tizoxanide enhances mtDNA biogenesis and mitophagy to shift heteroplasmy in MELAS cells. **A:** Oxygen consumption measured from MELAS cybrid cells treated with vehicle (DMSO) or tizoxanide (10 μM) for 48 h (n=4 biological replicates, error bars represent SE). **B:** HaeIII restriction enzyme digestion of a PCR-amplified fragment of mtDNA harboring the m.3243A>G mutation from DW7 cells treated with vehicle (DMSO) or tizoxanide (10 μM) for 144 h. The table shows n=4 biological replicates as mean ± SD. **C**: Genes identified using qPCR of MELAS cybrid cells stably overexpressing EV or MNRR1. **D:** MtDNA levels are shown relative to nuclear DNA (nDNA) (GAPDH) (n=5 biological replicates, error bars represent SE). **E:** Equal amounts of MELAS cybrid cell lysates, treated with vehicle (DMSO) or tizoxanide (10 μM) for 48 h, were separated on an SDS-PAGE gel and probed for PGC1α, PINK1, LC3 A/B, MNRR1, and Tubulin. **F:** Equal numbers of MELAS cells were treated as in **E**, separated on an SDS-PAGE gel, and probed for phospho-ubiquitin (S65) levels. Tubulin was probed as a loading control. In all figures, * indicates p<0.05, ** indicates p<0.005.

### MNRR1 activation using tizoxanide in MELAS patient fibroblasts enhances mitochondrial function and mitophagy and protects from LPS-induced inflammation

In primary fibroblasts from three independent MELAS patients, we found that activation of MNRR1 enhances mitophagy (as seen via LCB and pSer-65 ubiquitin levels) and mitochondrial biogenesis (PGC1⍺, MTCO2, TOM20 levels) (**Fig. 3A**). Furthermore, we found that OCR (**Fig. 3B**) and mitochondrial ATP (**Fig. 3C**) levels were enhanced. Since we recently found in a cell culture model that activation of MNRR1 resolved the effect of LPS-induced inflammation [16], and that mitochondrial diseases are associated with a pro-inflammatory phenotype [17], we also asked if we could rescue the effects of LPS in these patient fibroblasts. We found that LPS induces a pro-inflammatory response as judged by increased ROS and TNF levels, and this response can be blocked by tizoxanide (**Fig. 3D**), suggesting that MNRR1 activation is protective in these cells.

**Figure 3:**
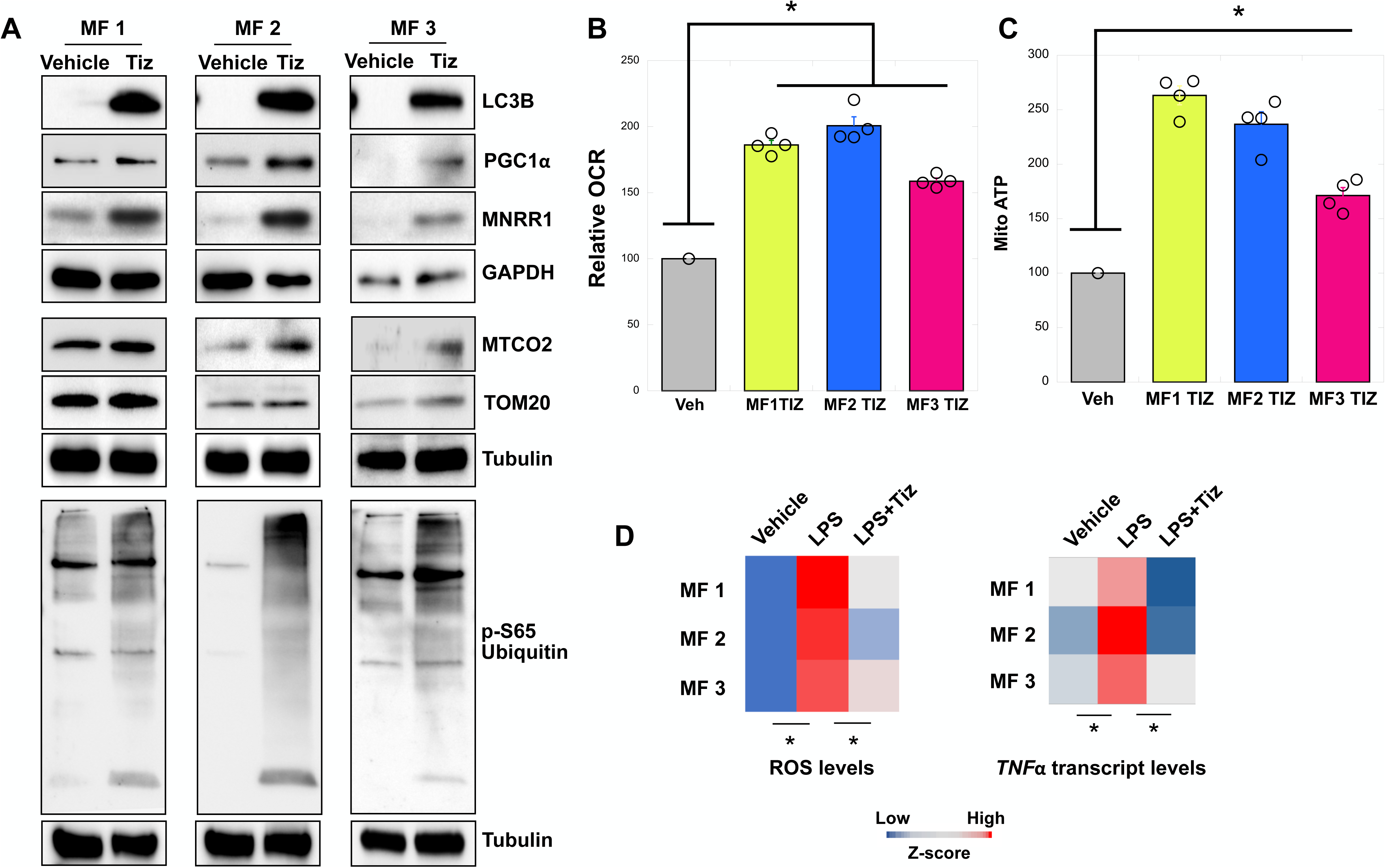
MNRR1 activation using tizoxanide enhances mitochondrial function, mitophagy, and protects from LPS-induced inflammation in MELAS patient fibroblasts. **A:** Equal amounts of lysates from MELAS patient fibroblast (MF 1, 2, or 3) lysates treated with vehicle (DMSO) or tizoxanide (10 μM) for 48 h were separated on an SDS-PAGE gel and probed for LC3B, PGC1α, MNRR1, MTCO2, TOM20, and phospho-ubiquitin (S65) levels. GAPDH or Tubulin was probed as a loading control. **B:** Oxygen consumption measured from MELAS patient fibroblasts treated with vehicle (DMSO) or tizoxanide (10 μM) for 48 h (n=4 biological replicates, mean ± SE). **C:** Mitochondrial ATP rate measured in MELAS patient fibroblasts treated with vehicle (DMSO) or tizoxanide (10 μM) for 48 h (n=4 biological replicates, mean ± SE). **D:** Heatmap representation of ROS and TNFa transcript levels from MELAS patient fibroblasts treated with vehicle (DMSO), LPS (DMSO + 500 ng/ mL LPS), or LPS plus tizoxanide (10 μM tizoxanide + 500 ng/mL LPS) for 24 h (n=4 biological replicates). In all figures, * indicates p<0.05, ** indicates p<0.005.

### Nitazoxanide acts by reducing HIF2α protein levels in MELAS cybrid cells

Nitazoxanide was identified by screening a library for transcriptional activators of *MNRR1* using 952-bp promoter luciferase-expressing stable cell lines. We therefore sought to identify the region on the MNRR1 promoter that responds to nitazoxanide with *MNRR1* induction. To this end we generated 200-bp deletions in the MNRR1 promoter and cloned them into the pGL4-basic luciferase vector (**Fig. 4A**). Upon testing the responsiveness of each of these constructs to nitazoxanide and tizoxanide in MELAS cybrid cells, we observed that a deletion of the 801-952 region on the promoter (D801-952) failed to display activation (**Fig. 4B**), suggesting that this promoter region was affected by nitazoxanide. Bioinformatic analysis of this region identified six bona fide binding sites for transcription factors (TFs) – Zeb1, HIF, ZFX, SMARCA3, ZNF35, and RBPJκ (**Fig. 4C**). RBPJκ binds to the core 13-bp element that we previously characterized to be responsive to moderate hypoxia and labeled as the oxygen responsive element (ORE) [11]. Of the six TFs, Picard et al. [6], who initially characterized the cells, identified only HIF2a to be transcriptionally induced (**Fig. 4D**). We assessed the level of both and found that HIF1α was not increased in the MELAS cells whereas HIF2α levels were higher in heteroplasmic MELAS cybrid cells (DW7) (**Fig. 4E**) as compared to the control cybrids (CL9) [6, 18]. To test whether tizoxanide was acting through HIF2α, we measured its protein levels and found that HIF2α was inversely proportional in a concentration dependent manner to MNRR1 in MELAS cybrid cells treated with tizoxanide (**Fig. 5A**). To test whether HIF2α is acting specifically via the hypoxia response element (HRE), we generated a deletion of the HRE in the MNRR1 promoter (**Fig. 4C**, **blue**). To our surprise, we found that the deletion of the HRE could not reverse the inhibition in MELAS cells, whereas deletion of the ORE could rescue the effects (**Fig. 5B**). To evaluate this confounding effect, we tested an HRE-harboring reporter in the MELAS cybrid cells (DW7) and found it to be more active than in the control cybrid cells (CL9) (**Fig. 5C**). In the same control cybrid cells, we could also overexpress HIF2α and repress MNRR1 transcription (**Fig. 5D**). However, since this effect was not through the HRE in the MNRR1 promoter (**Fig. 5B**), we again examined the sequence of the 800-952 region on the promoter and uncovered a second HRE in the reverse orientation on the opposite strand of the ORE where RBPJκ binds (**Fig. 5E**), thus providing a possible explanation for the effects seen in Figure 5B. We previously showed that MNRR1 forms a required transcriptional complex with RBPJκ at the ORE and that constitutively active RBPJκ can bypass the need for MNRR1 to activate transcription [11].

**Figure 4:**
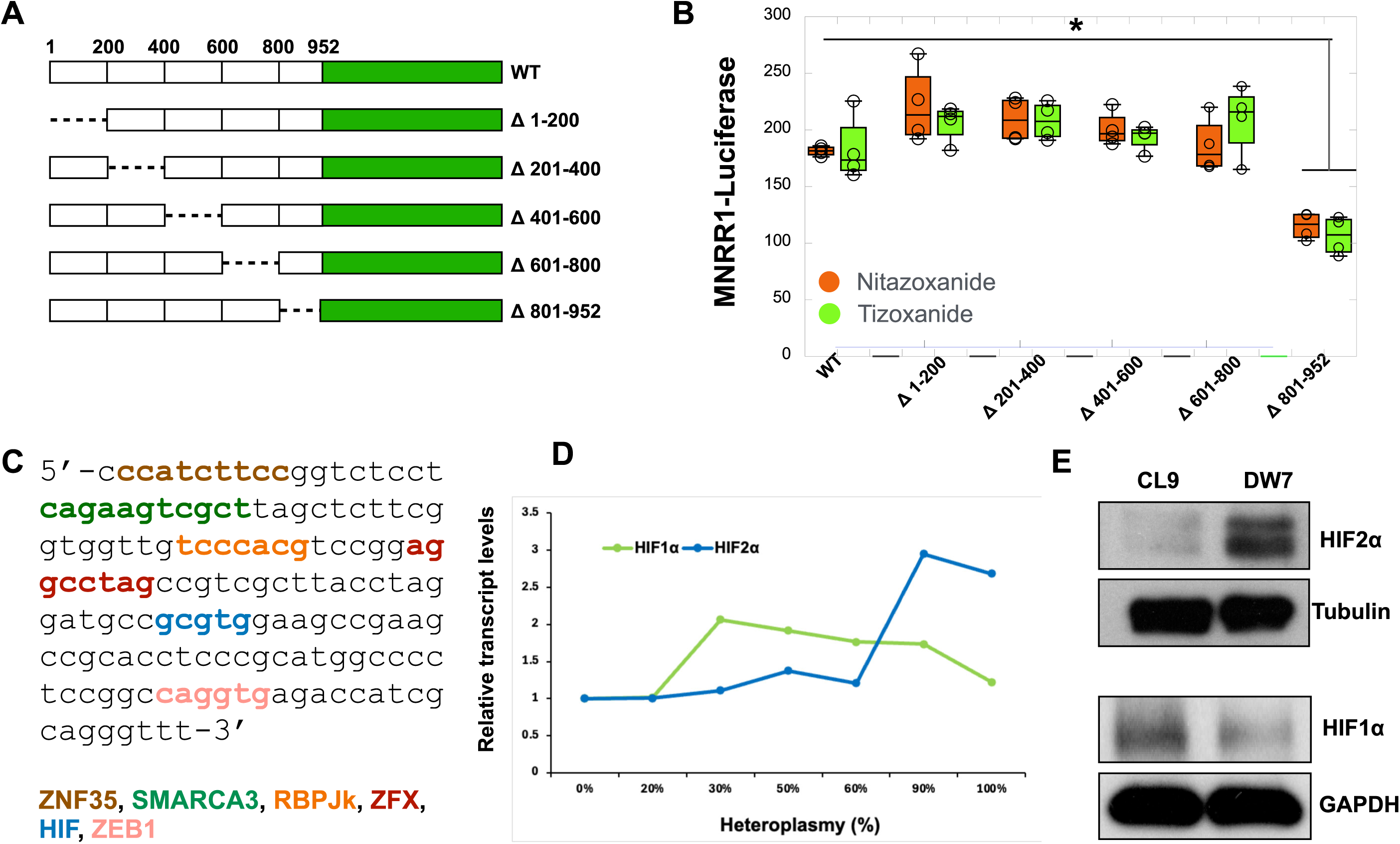
Nitazoxanide acts by reducing HIF2α levels in MELAS cells. **A:** Schematic representation of deleted regions of the MNRR1 promoter. **B:** Dual luciferase reporter assay using the MNRR1 promoter deletions in cells treated with vehicle (DMSO), nitazoxanide, or tizoxanide (10 μM) for 24 h (n=4 biological replicates). **C:** DNA sequence of region of 800-952 bp in the MNRR1 promoter highlighting binding sites of the transcription factors shown. **D:** Transcript levels of HIF2α and HIF1α in MELAS cybrid cells harboring different levels of heteroplasmy (0% to 100%). Data from [6]. **E:** Protein levels of HIF2α and HIF1α in cybrid cells with 0% (CL9) and 70% MELAS heteroplasmy (DW7) In all figures, * indicates p<0.05, ** indicates p<0.005.

**Figure 5:**
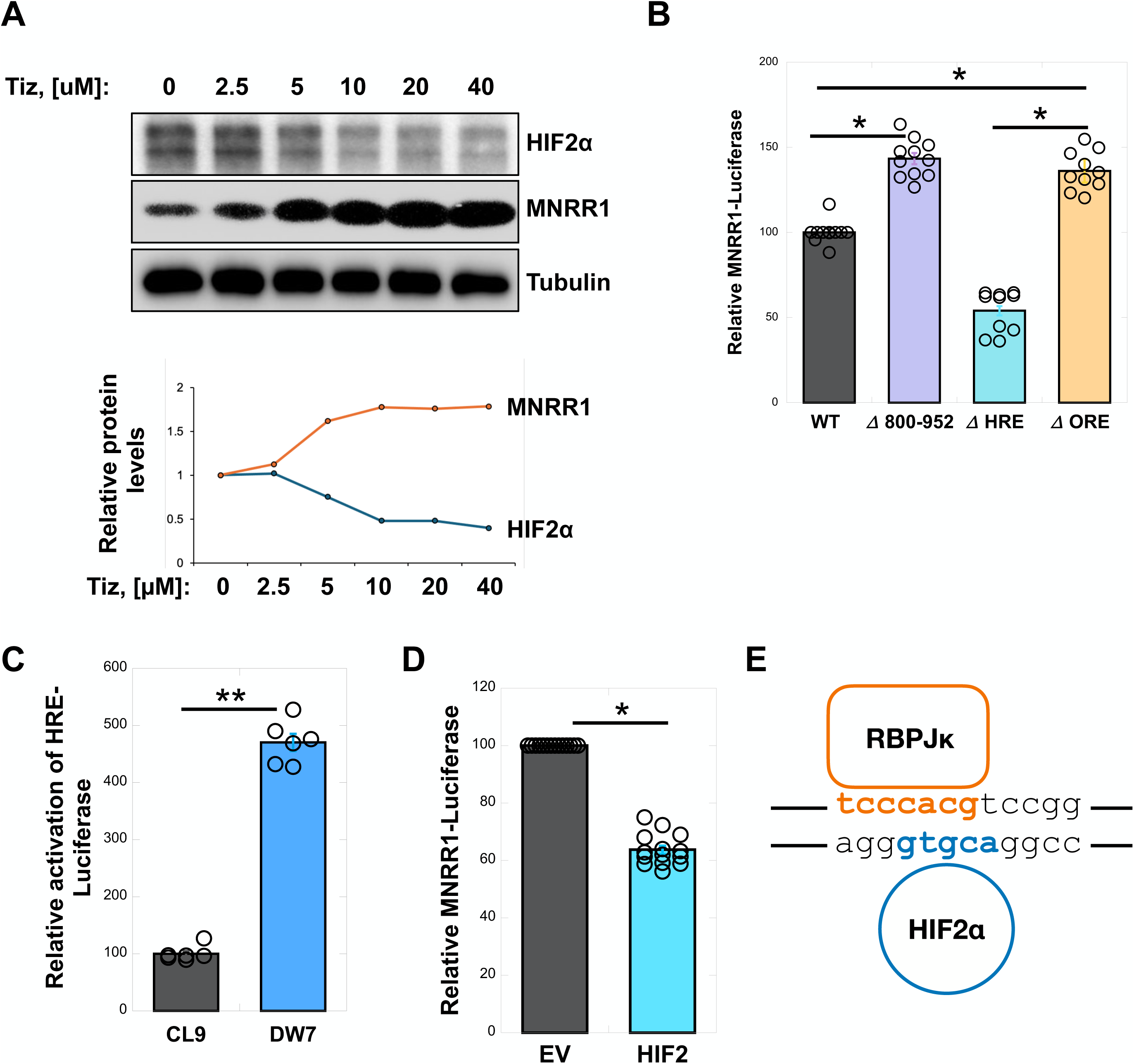
The reduction of MNRR1 in MELAS cells is via HIF2α acting at the ORE and not the HRE. **A:** *Above*, Equal amounts of MELAS cell lysates, treated with vehicle (DMSO) or increasing amounts of tizoxanide (2.5 to 40 μM) for 24 h, were separated on an SDS-PAGE gel and probed for HIF2α, MNRR1, and Tubulin. *Below,* Protein quantification for MNRR1 and HIF2α relative to Tubulin. **B:** Dual luciferase reporter assay showing relative activation of *MNRR1*-luciferase in MELAS cybrid cells overexpressing WT (Figure 4A), Δ 801-952 (Figure 4A), ΔORE (sequence in orange in Figure 4C) or ΔHRE (sequence in red in Figure 4C). **C:** Dual luciferase reporter assay showing relative activation of *HRE*-luciferase levels in cybrid cells with 0% (CL9) and 70% MELAS heteroplasmy (DW7). **D**: Dual luciferase reporter assay showing relative activation of *MNRR1*-luciferase levels in MELAS cybrid cells with 0% heteroplasmy (CL9) overexpressing EV (empty vector) or HIF2α. **E:** Sequences in *MNRR1* promoter highlighting the binding sites for RBPJκ (orange) and HIF2α (blue) on opposite strands of DNA. In all figures, * indicates p<0.05, ** indicates p<0.005.

### RBPJk and HIF2α compete for binding at the ORE in the MNRR1 promoter to regulate transcription

To dissect the effect of HIF2α and RBPJκ at the ORE, we first confirmed by chromatin immunoprecipitation that HIF2α can bind at the MNRR1 promoter (**Fig. 6A**). Using a constitutively active version of RBPJκ (CA-RBPJκ), we could rescue defective transcription of MNRR1 in MELAS cybrid cells (**Supp. Fig. 3A**) and block these effects by overexpression of HIF2 (**Supp. Fig. 3B**). We also titrated RBPJκ and HIF2α in MELAS cybrid cells and found that HIF2α can compete with, and inhibit, transcription induced by RBPJκ (**Fig. 6B**). Furthermore, a chemical inhibitor that blocks binding of RBPJκ to DNA (Auranonfin) was able to block the effects of MNRR1 induced transcription (**Supp. Fig. 3C**) whereas the effects of HIF2α overexpression were blocked by deletion of the ORE (**Fig. 6C**). Taken together with a previously published report that HIF interacts with RBPJk to inhibit transcription [19], we hypothesize that nitazoxanide relieves the inhibitory effect of HIF2α at the ORE to facilitate transcriptional activation via RBPJκ. At the protein level, depletion of HIF2α in the MELAS cybrid cells increases MNRR1 (**Fig. 6D**) and oxygen consumption (**Fig. 6E**). As a further confirmation, we specifically inhibited HIF2α with the compound PT2385 [20] and observed a rebound increase in both OCR and MNRR1 protein in MELAS cybrid cells (**Fig. 6F**). Consequently, we propose that HIF2α acts in two ways to regulate MNRR1 transcription: (1) it binds to the HIF site shown in blue (**Fig. 4C**) in the 800-952 bp region of the in the *MNRR1* promoter to activate transcription, and (2) it binds on the complementary stand of the ORE site shown in orange (**Fig. 4C**) to inhibit transcription. A balance between these mechanisms is imposed by the ratio of RBPJκ to HIF, as is suggested in Fig. 6B.

**Figure 6:**
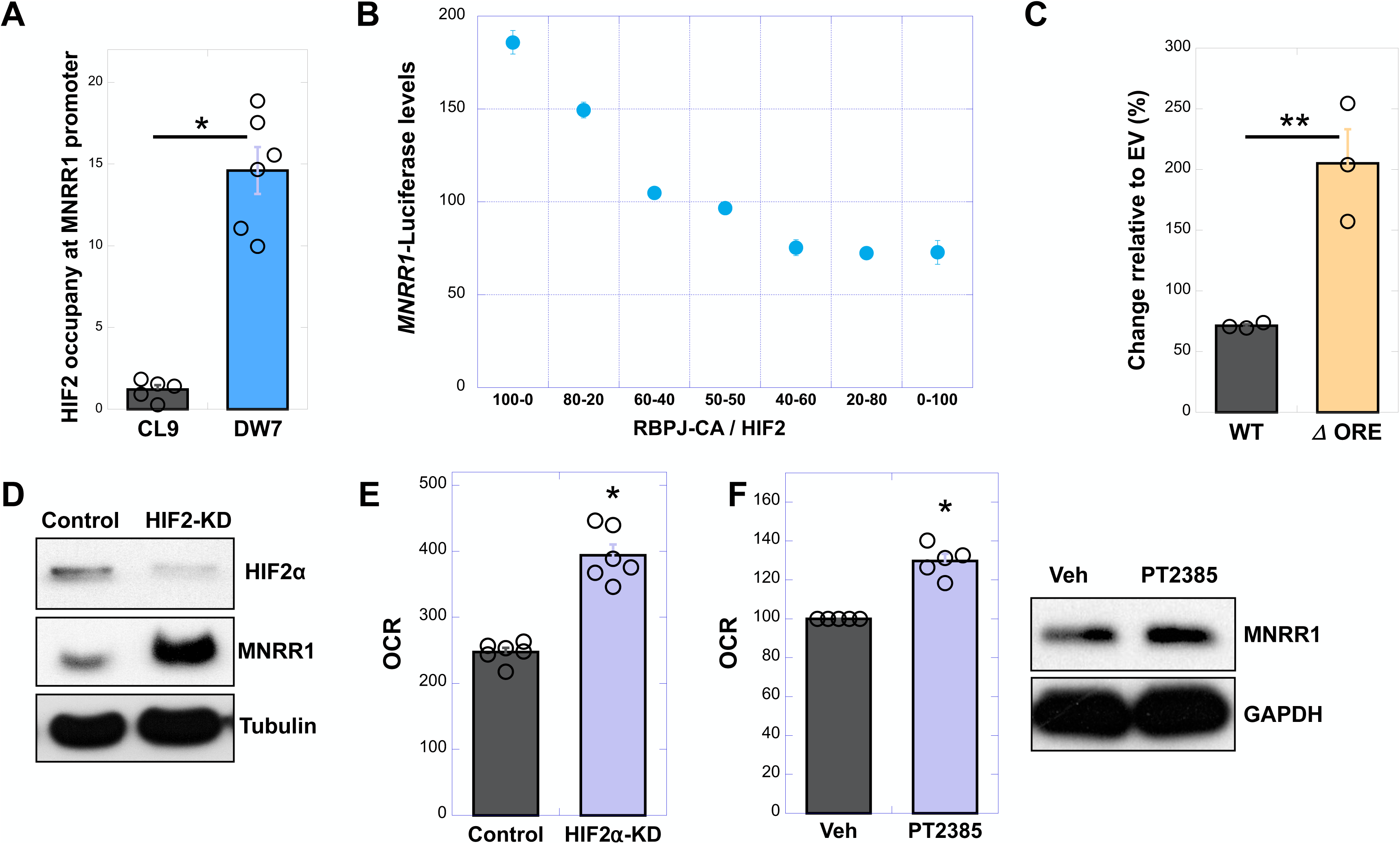
RBPJκ and HIF2α compete for binding at the ORE in the MNRR1 promoter to regulate transcription. **A:** Chromatin immunoprecipitation-qPCR assessing binding of HIF2α to the endogenous *MNRR1* promoter in MELAS cybrid cells with 0% (CL9) and 70% heteroplasmy (DW7). **B:** Dual luciferase reporter assay showing relative activation of *MNRR1*-luciferase levels in MELAS cybrid cells overexpressing varying proportions of constitutively active RBPJκ (RBP-CA) and HIF2α. **C:** Dual luciferase reporter assay showing relative activation of *MNRR1*-luciferase levels in MELAS cybrid cells overexpressing the WT (Figure 4A) or ΔORE (sequence in orange in Figure 4C) promoter with EV (dotted line) or HIF2α. **D:** Equal amounts of Control or HIFα knockdown (KD) MELAS cybrid cell lysates were separated on an SDS-PAGE gel and probed for HIF2α and MNRR1 levels. Tubulin was probed as loading control. **E:** Oxygen consumption measured from Control or HIFα knockdown (KD) MELAS cybrid cells. (n=4 biological replicates, error bars represent SE). **F:** *Left,* Oxygen consumption measured from MELAS cybrid cells treated with vehicle (DMSO) or PT2385 (10 μM), an inhibitor of HIF2α function, for 24 h. *Right*, Equal amounts of MELAS cybrid cell lysates treated with vehicle (DMSO) or PT2385 (10 μM) for 24 h, separated on an SDS-PAGE gel, and probed for MNRR1 levels. GAPDH was probed as a loading control.

### PHD3 levels are reduced in MELAS cybrid cells and enhanced by nitazoxanide to increase MNRR1 levels

The absence of any change in HIF2α transcript levels with nitazoxanide and tizoxanide (**Supplementary Fig. 4**) suggested that HIF was showing greater protein stability. Hence, we assessed the levels of all three prolyl hydroxylases (PHD1, 2, and 3) in the MELAS cybrid cells using published transcriptomics data (**Fig. 7A**) [6]. Of these, we found that only PHD2, which shows specificity for HIF1α, was increased at high heteroplasmy, consistent with the reduction of HIF1α in DW7 versus CL9 cells (**Fig. 4E**). Next, we assessed the levels of PHD3 and found that it was reduced in the DW7 MELAS cells (**Fig. 7B**). Since HIF2α is increased in DW7 cells (**Fig. 4E**), we examined the role of PHD1 and PHD3. We found that PHD3 is most increased by tizoxanide at the protein level (**Fig. 7C**) and, when overexpressed, uniquely stimulates respiration in DW7 cells (**Fig. 7D**), suggesting that PHD3 may be the prolyl hydroxylase responsible for targeting HIF2α. PHD2 was not evaluated here as studies have shown this enzyme to be selective for HIF1 [21, 22], whose levels in MELAS cells were not correlated with heteroplasmy [6]. These results suggest that nitazoxanide and tizoxanide transcriptionally induce *MNRR1* by reducing the inhibitory effect of HIF2α and facilitating its activation via RBPJκ.

**Figure 7:**
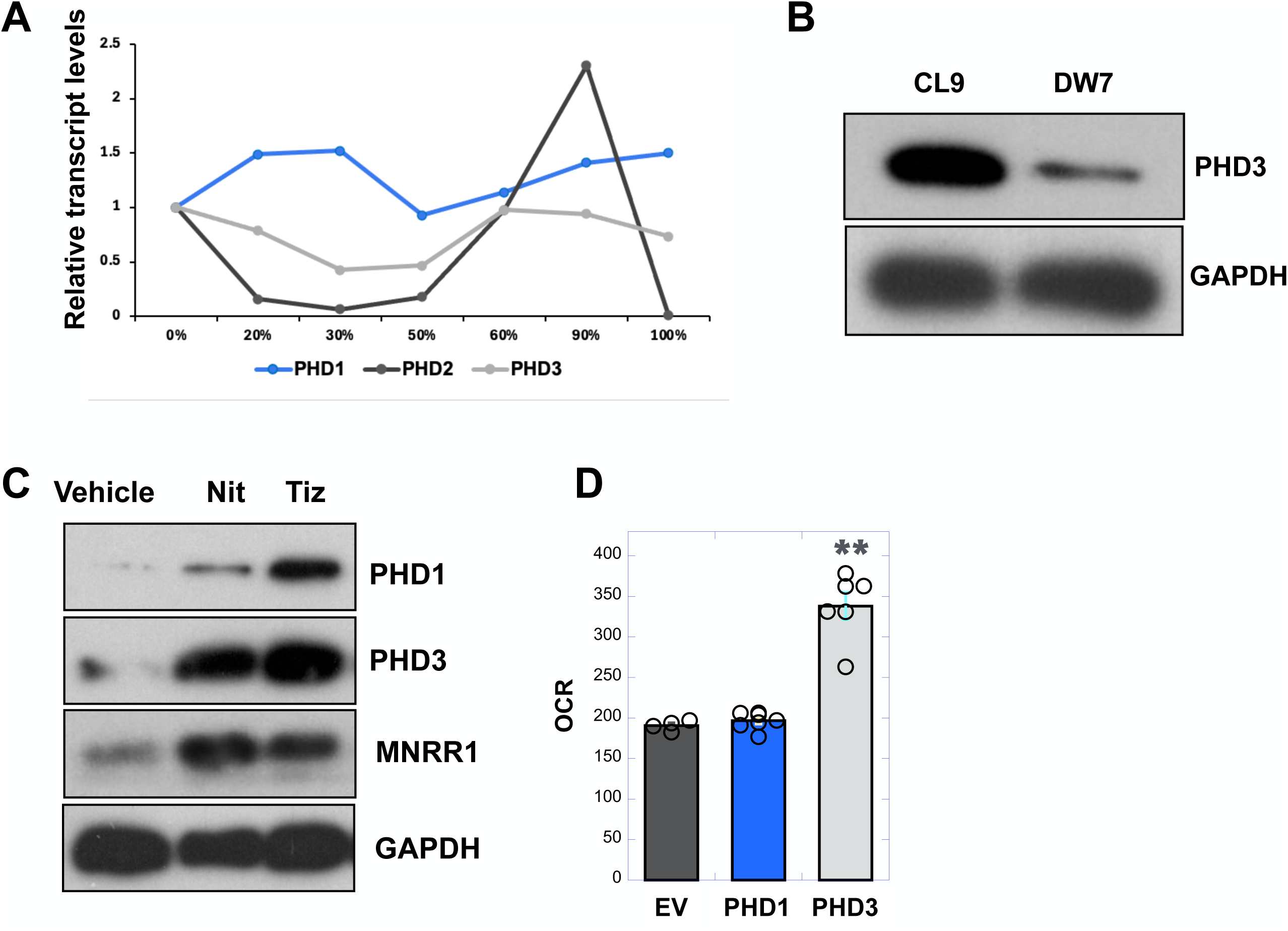
PHD3 levels are reduced in MELAS cybrid cells and enhanced by nitazoxanide to increase MNRR1 levels. **A:** Transcript levels of PHD1, 2, and 3 in MELAS cybrid cells harboring the levels of heteroplasmy shown (data from [6]). **B:** Protein levels of PHD3 in MELAS cybrid cells with 0% (CL9) and 70% heteroplasmy (DW7). GAPDH was probed as loading control. **C:** Equal amounts of MELAS cybrid cell lysates, treated with vehicle (DMSO), nitazoxanide, or tizoxanide (10 μM) for 24h, were separated on an SDS-PAGE gel and probed for PHD1, PHD3, MNRR1, and Tubulin. **D:** Oxygen consumption measured from MELAS cybrid cells overexpressing PHD1 or PHD3. (n=4 biological replicates, error bars show SE).

## Discussion

We previously showed that MELAS cells contain a reduced amount of MNRR1 and that genetically restoring the amount alleviated much of the pathophysiological phenotype such as reduced energy generation and increased ROS [12]. This suggested that activation of MNRR1 could improve mitochondrial deficits associated with MELAS and stimulated a search for a small molecule that would restore MNRR1 levels. We discovered the MNRR1 transcriptional activator nitazoxanide by high throughput screening of a drug and natural products chemical library. We found that it and its metabolic breakdown product tizoxanide can restore expression and thereby function in MELAS cybrid cells and, importantly, in primary fibroblasts from MELAS patients. In seeking to identify the drug’s mechanism of action, we uncovered that MELAS cells contain increased amounts of HIF2 at normoxia, and that HIF2 binds at the *MNRR1* promoter to inhibit transcription.

In parasites nitazoxanide inhibits pyruvate:ferredoxin oxidoreductase (PFOR), a key enzyme utilized by anaerobes in the oxidative decarboxylation of pyruvate to acetyl-CoA and CO_2_, whereas in humans nitazoxanide has no known target. We first identified the region of the MNRR1 promoter where nitazoxanide acts and then narrowed to the specific transcription factors – HIF2α and RBPJκ – that bind to overlapping regions in the ORE part of the promoter to regulate transcription (**Figure 5E**). We have previously shown that RBPJκ binds to and mediates transcription at the ORE [11] and now identify a second factor – HIF2α – that can bind to the MNRR1 promoter (**Figure 5E**) to regulate transcription. These factors compete with each other: active RBPJκ can enhance transcription of MNRR1 and HIF2α competes with and represses this effect. In MELAS cybrid cells, which show a pseudohypoxic state, expressing higher levels of HIF2α, MNRR1 is transcriptionally inhibited. With regard to the action of nitazoxanide, we propose a novel mechanism of post-translational regulation: we found that PHD3, the prolyl hydroxylase that is largely responsible for targeting HIF2α for degradation by the ubiquitin proteasome [23, 24], is upregulated by tizoxanide, thereby increasing MNRR1 levels. Since the anti-inflammatory effect of nitazoxanide was explicitly shown to depend on MNRR1 [13], this effect of nitazoxanide will be an important focus of future studies.

The patient fibroblasts do not show a shift of heteroplasmy after tizoxanide treatment (data not shown), unlike the shift toward wild-type seen in 70% MELAS cybrid cells (**Figure 2B**). This may result from the low heteroplasmy levels (<50%) in the patient fibroblasts, consistent with distinct nuclear responses for each heteroplasmy range previously seen in the cybrid model of MELAS [6]. Lower heteroplasmy levels may not induce sufficient stress in these glucose-grown cells to alter heteroplasmy.

Nitazoxanide is the prodrug formulation of tizoxanide, a commonly used FDA-approved antiprotozoal effective against diarrheal symptoms caused by *Giardia* or *Cryptosporidium*. Nitazoxanide is deacetylated *in vivo* to tizoxanide, which has antioxidant properties and is a known inhibitor of iNOS [25]. Both properties are likely major contributors for the observed improvements in lung and other organ damage in SARS-Cov-2 patients [26]. Consistently, MNRR1 reduction increases ROS production [9]. Thus, nitazoxanide has the potential to also alleviate MELAS beyond activation of MNRR1 transcription. Furthermore, this recently rekindled interest in the broad spectrum of tizoxanide applications to human health has led to new inquiry. The generation of more plasma-stable congeners that replace the acetyl group substituent attached to the hydroxyl group with a more stable formamyl group [27], as well as novel approaches for tizoxanide quantification *in vitro* and *in vivo*, have recently been published [28, 29]. Further studies will elucidate how increased plasma concentrations and systemic exposure to the active tizoxanide moiety will be tolerated and distributed in *in vivo* models of infection, cancer, and other maladies with unmet needs.

In summary, we have identified nitazoxanide and its metabolite tizoxanide as a drug that stimulates MNRR1 transcription and shown that it reduces heteroplasmy in a cell culture model of MELAS and improves the phenotype in MELAS patient fibroblasts. Given its established safety profile, it could be usefully evaluated for diseases like MELAS and other inflammatory conditions wherein MNRR1 levels become reduced [16]. The observation that activation of MNRR1 can protect from inflammatory stress is crucial since patients with MELAS and other mitochondrial diseases have a higher susceptibility to infections and bacterial sepsis [17]. Hence, these results are also consistent with our recent findings that activation of MNRR1 can prevent inflammation induced preterm birth *in vivo* [13]. Furthermore, we have identified increased HIF2 at normoxia as the cause of reduced MNRR1 in MELAS cells, uncovering a potential new therapeutic target. By contrast, most current studies target symptomatic relief such as by increasing nitric oxide levels to ameliorate episodic stroke-like episodes [*e.g.* 26].

## Materials and Methods

### Cell lines

The human embryonic kidney cell line HEK293, the triple-negative breast cancer cell line MDA-MB-468, the human first trimester placental cells HTR8/SVNeo (HTR), SHSY5Y cells, and HMC3 were obtained from the ATCC (Manassas, VA). The MELAS cells (143B human ostersarcoma cybrid) were a kind gift from Dr. Douglas Wallace. The HEK293 and human fibroblast cells were cultured in DMEM with L-glutamine and D-Glucose (Gibco, Billings, MO) supplemented with penicillin-streptomycin (HyClone, Logan, MT) and 10% fetal bovine serum (FBS) (Sigma Aldrich, St. Louis, MO); the MDA-MB-468 cells were cultured as above but plus 1 mM pyruvate; the HTR were cultured in Roswell Park Memorial Institute Medium (RPMI) (HyClone) supplemented with 5% FBS plus penicillin-streptomycin; and the MELAS cells were grown in DMEM with 1 mM pyruvate supplemented with non-essential amino acids (Gibco), 50 μg/ml uridine, and 10% FBS. The SHSY5Y cells were cultured DMEM/F12 with L-glutamine and D-Glucose (Corning, Corning, NY) supplemented with Penicillin-Streptomycin (HyClone) and 10% fetal bovine serum (FBS) (Sigma Aldrich) in EMEM with the same additives.

### Chemicals

Tizoxanide, nitazoxanide, and genetisate were obtained from Selleckchem (Houston, TX), Phenyl-4-aminosalicylic acid was obtained from A2Bchem (San Diego, CA), 2-Amino 5-nitrothiazole was from Sigma, and the remaining compounds (4-Aminobenzanilide, 4-Amino salicylic acid, benzanilide, phenyl benzoate, and phenyl salicylic acid) were from Santa Cruz Biotechnology (Dallas, TX). All compounds were solubilized in DMSO (used as vehicle control in all experiments with these compounds). LPS (Lipopolysaccharide from *Escherichia coli* 0111:B4) was purchased from Invivogen.

### Plasmids

The *MNRR1* promoter luciferase reporter plasmid and pRL-SV40 *Renilla* luciferase expression plasmids have been described previously [11]. The HRE-*Luciferase* plasmid was purchased from Addgene (#26731). All expression plasmids were purified using the EndoFree plasmid purification kit from Qiagen (Germantown, MD).

### Cell lines, culture, and Z’ determination

#### Stable cell line generation

HEK293 or MDA-MB-468 cells were transfected with pGL4-MNNR1-luciferase, selected with 0.5 μg/mL (for 293) or 1 μg/mL (for 468) puromycin for 2 weeks, and subcloned by limiting dilution. MDA-MB-468-MNRR1-luc and HEK293-MNRR1-luc cells from multiple clones (5000 -7500 cells) were plated in 100 μL of complete medium overnight and treated with 10 μM of each compound from the library for 24 h. The medium was aspirated from each well to a volume of 25 μL, and 25 μL Bright Glo (Promega, Madison, WI) luciferase detection reagent was added to each well 10 min prior to luminescent determination with a FlexStation 3 Multimode Microplate Reader (Molecular Devices, San Jose, CA). In the absence of a known MNRR1-inducing small molecule, we chose MNRR1 overexpression (since MNRR1 induces its own expression [9]) as a potential positive control. The Z’ factor is a value used to identify strong candidates and can be calculated using a previously described method [30]. The range of this value is negative infinity to one, with > 0.5 as a very good assay, > 0 an acceptable assay, and < 0 an unacceptable assay. The values obtained for this screen were >0, but we were unable to identify any candidates with strong values of 0.5 to 1.

#### MicroSource Spectrum Collection

Despite the low Z’ values for MDA-MB-468-MNRR1-luc and HEK293-MNRR1-luc, pilot screens were performed with both using the MicroSource Spectrum Collection (Gaylordsville, CT). The Spectrum Collection is comprised of ∼2400 small molecules and natural products that are known drugs or otherwise biologically well-characterized. This library contains a manageable number of compounds in 10 mM DMSO stocks that can be tested without the need for advanced liquid handling. Additionally, the use of biologically well-characterized compounds facilitates the rapid identification of pathways and signaling networks likely to be of interest to the investigator. Dry powder stocks of the compounds that provided the most robust response in both cell lines and that did not have chemical liabilities that would preclude their use in cultured cells or, potentially, in human subjects, were then obtained from commercial sources. For HTS, cells were plated, treated, and measured for luciferase expression as described for clone identification; all HTS compounds were added to a final concentration of 10 μM and MNRR1-luciferase expression was measured after 24 h.

Since highly expressing clones that provided a Z’ value between 0.5 – 1.0 were elusive and there is a general lack of MNRR1 activators that could be used as positive controls, the criterion of accepting modestly enhanced transcription (1.8 – 2.8-fold) in each of two cell lines was used. Thus, a pilot screen with the MicroSource Spectrum Collection was performed in MNRR1-luciferase clones from MDA-MB-468-*MNRR1*-luc and HEK-293-*MNRR1*-luc cells. Compounds enhancing transcription in both cell lines were considered for further scrutiny with orthogonal assays to evaluate *MNRR1* gene and protein expression. Nitazoxanide emerged as a validated MNRR1 activator and was studied further using *in vitro* assays in MELAS cell lines and primary fibroblasts to determine whether chemically induced MNRR1 expression could improve known human pathologic mitochondrial deficiencies. In addition to the lack of a known positive control, we hypothesize that the low Z’ values observed with our *MNRR1*-luciferase cell lines were due to the relatively low levels of MNRR1 expressed at baseline in monolayer cultures of MDA-MB-468 and HEK293.

### Transient transfection of MELAS cells

MELAS cells were transfected with the indicated plasmids using TransFast transfection reagent (Promega) according to the manufacturer’s protocol. A TransFast:DNA ratio of 3:1 in serum and antibiotic free medium was used. Following incubation at room temperature for ∼15 min, the cells were overlaid with the mixture. The plates were incubated for 1 h at 37 °C followed by replacement with complete medium and further incubation for the indicated time.

### Real-time polymerase chain reaction (RT-PCR)

Total cellular RNA was extracted from MELAS cells with a RNeasy Plus Mini Kit (Qiagen) according to the manufacturer’s instructions. Complementary DNA (cDNA) was generated by reverse transcriptase polymerase chain reaction (PCR) using the ProtoScript® II First Strand cDNA Synthesis Kit (New England Biolabs, Ipswich, MA). Transcript levels were measured by real time PCR using SYBR green on an ABI 7500 system. Real-time analysis was performed by the ΔΔ^Ct^ method [31]. The primers used were *MNRR1* forward: 5′-CACACATGGGTCACGCCATTACT-3′, reverse: 5′-TTCTGGGCACACTCCAGAAACTGT-3′; 18s forward: 5′-CCAGTAAGTGCGGGTCATAA-3′, reverse: 5′-GGCCTCACTAAACCATCCAA-3′, *PHD1*(*EGLN2)* forward: 5′-ACATCGAGCCACTCTTTGAC-3’, reverse: 3’-TCCTTGGCATCAAAATACC-5’[32]; *PHD3(EGLN3)* forward 5’-TCAAGGAGAGGTCTAAGGCAA-3’, reverse: 3’-ATGCAGGTGATGCAGCGA-5’ [33] and *HIF2⍺ (EPAS1)*forward: 5′-CACCAAGGGTCAGGTAGTAA-3’, reverse: 3′-AACACCACGTCATTCTTCTC-5’.

### Luciferase reporter assay

Luciferase assays were performed with the dual-luciferase reporter assay kit (Promega). Briefly, cells were lysed in 1x passive lysis buffer (Promega) and 25 μL of lysate was used for assay with a tube luminometer using an integration time of 10 s. Transfection efficiency was normalized with the co-transfected pRL-SV40 *Renilla* luciferase expression plasmid [8, 11].

### Immunoblotting

Immunoblotting was performed as described previously [11, 31]. Cell lysates for immunoblotting were prepared using RIPA buffer (Abcam, Waltham, MA) and included a protease and phosphatase inhibitor cocktail (Sigma, St. Louis, MO). Total protein extracts were obtained by centrifugation at 21,000 x *g* for 30 min at 4 °C. The clear supernatants were transferred to new tubes and quantified using the Bradford reagent with BSA as standard (BioRad, Hercules. CA). Equal amounts of cell lysates were separated by sodium dodecyl sulphate–polyacrylamide gel electrophoresis (SDS–PAGE), transferred to PVDF membranes (BioRad), and blocked with 5% non-fat dry milk. Incubation with primary antibodies (used at a concentration of 1:500) was performed overnight at 4 °C. The PGC1α (catalog no. 2178), PINK1 (6946), LC3A/B (12741), phosphoserine-65 ubiquitin (62802), HIF2α (59973), TOM20 (72610), GAPDH (8884), actin (12748), and tubulin (9099) antibodies were obtained from Cell Signaling (Danvers, MA). The MNRR1 (19424-1), MTCO2 (55070-1), PHD1 (12984-1) and PHD3 (18325-1) antibodies were obtained from Proteintech (Chicago, IL). Incubation with secondary antibodies (1:5000) was performed for 2 h at room temperature. For detection after immunoblotting, the SuperSignal™ West Pico PLUS substrate (ThermoFisher, Waltham, MA) was used to generate chemiluminescence signal, which was detected with X-Ray film (RadTech, Vassar, MI).

### Immunofluorescence

Cells plated on glass cover slips were fixed with 3.7% formaldehyde (prepared in 1x PBS) at room temperature for 15 min, followed by permeabilization with 0.15% Triton X-100 (prepared in distilled water) for 2 min, and then blocked with 5% bovine serum albumin (BSA) (prepared in 1x PBS, 0.1% TWEEN-20 (PBST)) for 1 h at room temperature. Cells were washed with PBST then incubated for 1 h at room temperature in primary antibody solution containing Coralite® 594 conjugated mouse monoclonal anti-CHCHD2 IgG (1:100, Proteintech, Cat. No. CL594-66302) prepared in PBST. Cells were washed 3 times with PBST for 5 min each and mounted with Vectashield vibrance with DAPI (Cat. # H-1800-10, Vector Labs, Newark, CA). Cells were imaged at 63x on the confocal 60 μm disk setting with the BioTek Cytation C10 using the Gen5 software (Agilent). Six fields for each group were taken and z-stacks of 25 slices (±3 slices) were performed for each field followed by a z-projection and image deconvolution. Corrected total fluorescence for each field to determine MNRR1 content was calculated using FIJI (National Institutes of Health).

### Mitochondrial DNA Levels

Total genomic DNA was isolated from cells expressing each of the mutants using the Invitrogen PureLink Genomic DNA Mini Kit (Thermo Fisher Scientific, K1820-01) and analyzed by real-time PCR as above. The primer sequences used to amplify mtDNA and GAPDH were as follows: mtDNA forward: 5′-CCTCCCTGTACGAAAGGAC-3′; reverse: 5′-GCGATTAGAATGGGTACAATG-3′; GAPDH forward: 5′-GAGTCAACGGATTTGGTCGT-3′; reverse: 5′-TTGATTTTGGAGGGATCTCG -3′.

### Restriction Enzyme Digestion

DNA was analyzed for the MELAS mutation as described previously [12]. The A→G mutation creates a new HaeIII site at position 3243 that can be amplified by PCR using primers corresponding to the light-strand positions 3116 to 3134 and to the heavy-strand positions 3353 to 3333. Equal amounts of the resulting products were digested with the restriction enzyme HaeIII (New England Biolabs) and electrophoresed on a 2.5% agarose gel.

### ROS measurements

Total cellular ROS measurements were performed with CM-H_2_DCFDA (Life Technologies). Cells were distributed into 96-well plates at 2.5 x 10^4^ cells per well and incubated for 24 h or as described in specific experiments. Cells were then treated with 10 μM CM-H_2_DCFDA in serum-and antibiotic-free medium for 1 h. Cells were washed twice in phosphate buffered saline and analyzed for fluorescence on a BioTek Synergy H1 Microplate Reader (Agilent).

### Intact cellular oxygen consumption and measurement of ATP

Cellular oxygen consumption was measured with a Seahorse XF^e^24 Bioanalyzer (Agilent). Cells were plated at a concentration of 3.5 x 10^4^ per well a day prior to treatment and basal oxygen consumption was measured 48 h after treatments, as described [8, 31]. For ATP levels, Agilent Seahorse ATP Real-Time rate assay kit was used per manufacturer’s instructions.

### Chromatin Immunoprecipitation-qPCR (ChIP-qPCR)

Chromatin immunoprecipitation was performed per manufacturer’s instructions using SimpleChIP® Enzymatic Chromatin IP Kit (Cell Signaling, 9002). Briefly, 2x10^7^ cells were fixed with formaldehyde to crosslink and chromatin was digested into ∼150-900bp fragments using a combination of micrococcal nuclease and sonication. 2% of the sample was stored as input control. This digested chromatin was immunoprecipiated using the using the HIF2α antibody (Cell Signaling, 59973). Samples were eluted and the crosslinking was reversed. The eluted DNA and the input controls were purified and tested for relative amplification using qPCR analysis. The primers used were as follows: forward: 5’-ATCTTCCGGTCTCCTCAGAA-3’; reverse: 3’-AAACCCTGCGATGGTCTCA-5’.

### Statistical Analysis

All statistical analyses were performed with the two-sided Wilcoxon rank sum test using MSTAT version 6.1.1 (N. Drinkwater, University of Wisconsin–Madison). *P < 0.05; **P < 0.005.

## Acknowledgements

We thank Douglas Wallace, University of Pennsylvania, for providing cybrid cells containing the MELAS mtDNA mutation.

## Data availability

All study data presented in this manuscript are included in the article or are available from the lead contacts upon request. Unique reagents generated from this study are available from the lead contacts with a completed Materials Transfer Agreement. This study did not generate original code.

---

**Supplementary Figure 1:**
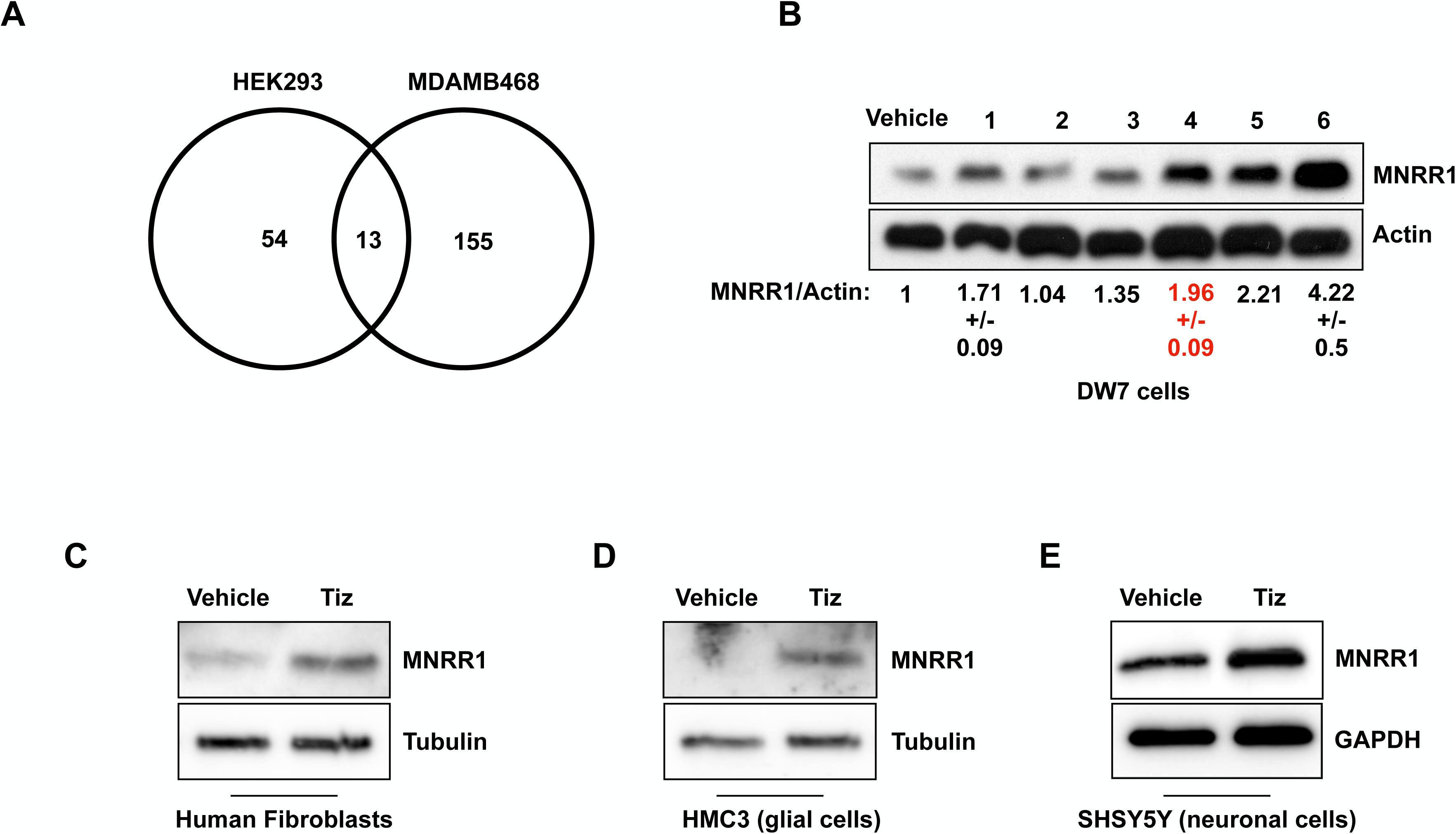
**A:** Venn diagram showing activators identifed in HEK293 and MDA-MB-468 cells. **B:** Equal amounts of MELAS cells treated with Vehicle (DMSO) or various MNRR1 activating compounds (10 μM) for 24 h were separated on an SDS-PAGE gel and probed for MNRR1 levels. Actin was probed as a loading control and numbers below represent an average and standard deviation (SD) of two biological replicates. **C-E:** Equal amounts of cell lysates from various cell lines treated with vehicle (DMSO) or tizoxanide (10 μM) for 24 h were separated on an SDS-PAGE gel and probed for MNRR1 plus loading controls GAPDH or tubulin.

**Supplementary Figure 2:**
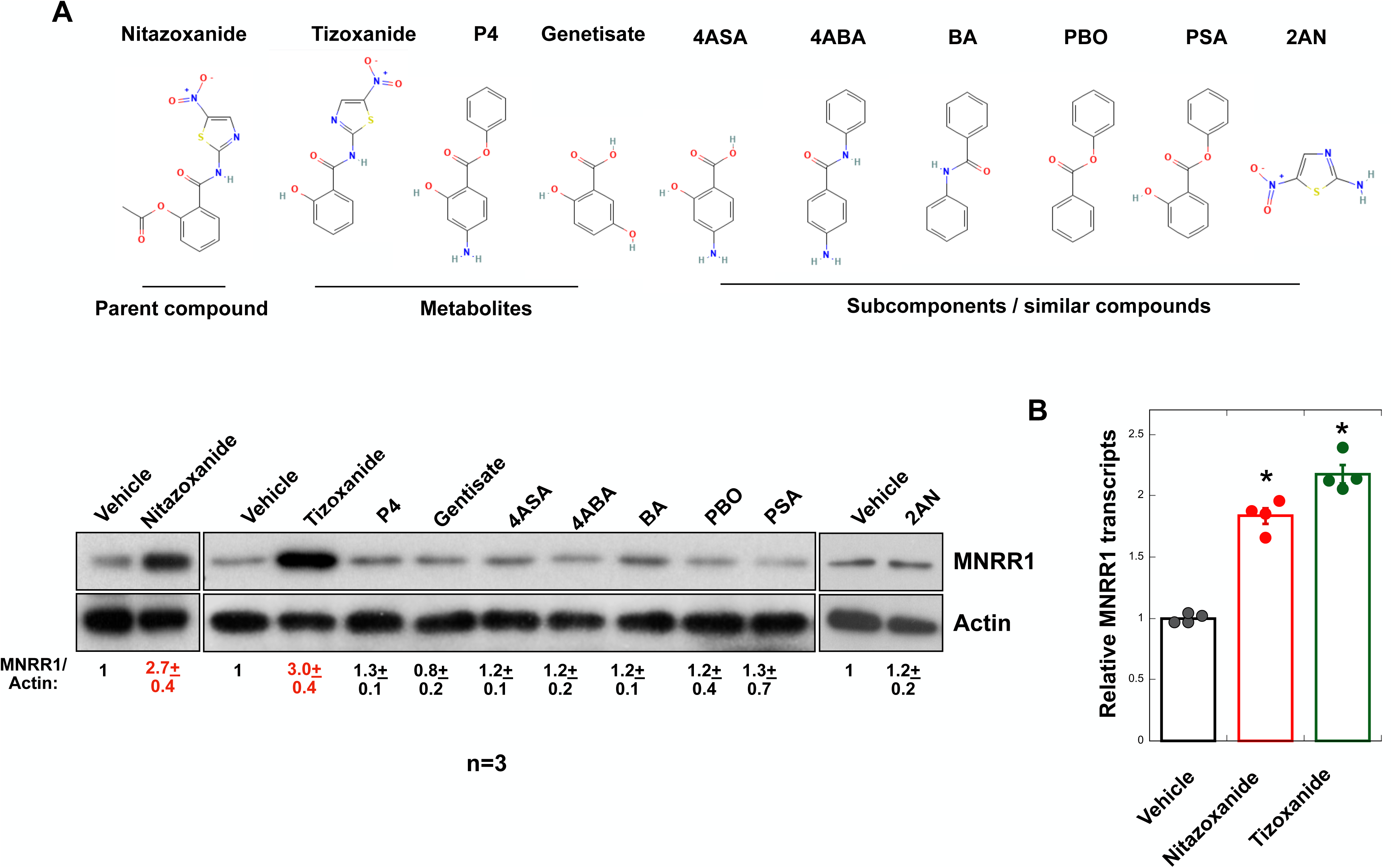
**A:** *Above*, Chemical structures of nitazoxanide, its metabolites, and similar compounds. Abbreviations: P4, Phenyl-4-aminosalicylic acid; 4ABA, 4-Aminobenzanilide; 4ASA, 4-Amino salicylic acid; BA, Benzanilide; PBO, Phenyl benzoate; PSA, Phenyl salicylic acid; 2AN, 2-Amino 5-nitrothiazole. *Below*, Equal amounts of lysates of MELAS cells treated with Vehicle (DMSO) or the various compounds (10 μM) for 24 h were separated on an SDS-PAGE gel and probed for MNRR1 levels. Actin was probed as a loading control and numbers below represent an average and SD of 3 biological replicates. **B:** *MNRR1* transcript levels are shown relative to 18S rRNA (n=4 biological replicates, error bars represent SE). In all figures, * indicates p<0.05, ** indicates p<0.005.

**Supplementary Figure 3:**
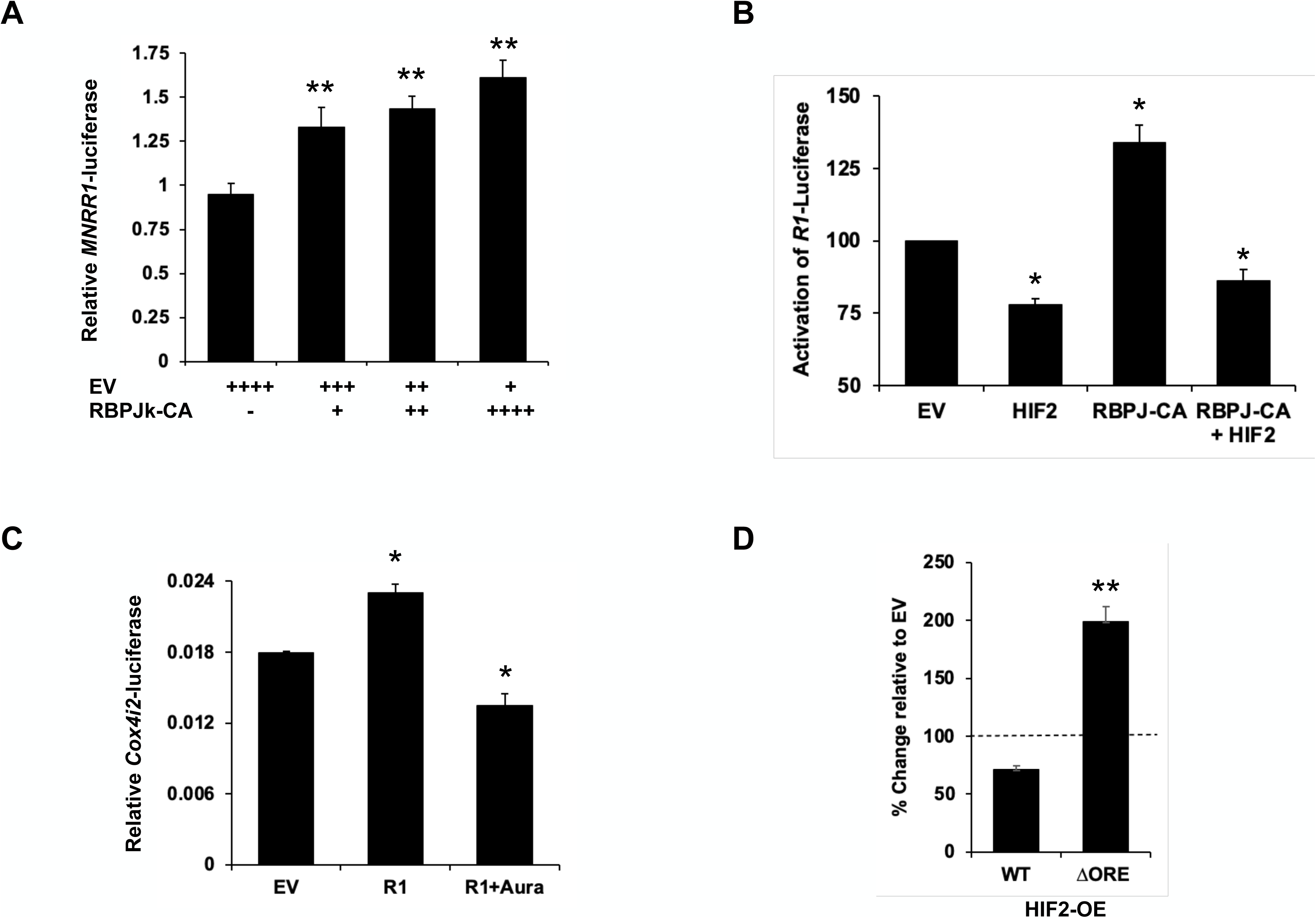
**A:** Dual luciferase reporter assay showing relative activation of *MNRR1*-luciferase levels in MELAS cybrid cells overexpressing varying proportions of empty vector (EV) and constitutively active RBPJκ (RBP-CA). **B:** Dual luciferase reporter assay showing relative activation of *MNRR1*-luciferase levels in MELAS cybrid cells overexpressing an empty vector (EV) or constitutively active RBPJκ (RBP-CA) and HIF2α. **C:** Dual luciferase reporter assay showing relative activation of *COX4I2*-luciferase levels in MELAS cybrid cells overexpressing an empty vector (EV), MNRR1, and MNRR1+Auranofin (0.5μM).

**Supplementary Figure 4:**
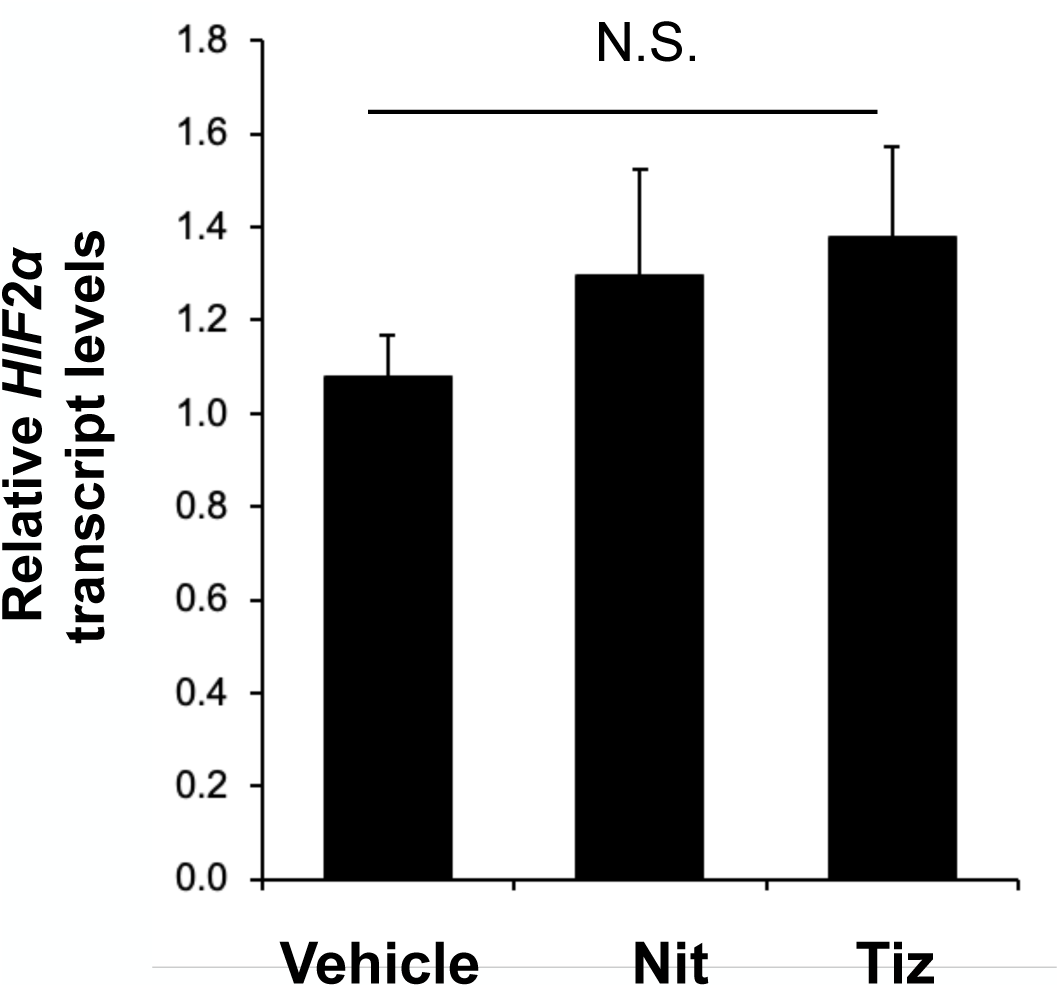
RT-PCR for measuring *HIF2α* levels. 18S rRNA was used as housekeeper for both analyses (n=3 biological replicates).

## References

1. Nass, S. and M.M. Nass, Intramitochondrial Fibers with DNA Characteristics. Ii. Enzymatic and Other Hydrolytic Treatments. J Cell Biol, 1963. 19(3): p. 613–29.

2. Nass, M.M. and S. Nass, Intramitochondrial Fibers with DNA Characteristics. I. Fixation and Electron Staining Reactions. J Cell Biol, 1963. 19(3): p. 593–611.

3. Chalmers, R.M. and A.H. Schapira, Clinical, biochemical and molecular genetic features of Leber’s hereditary optic neuropathy. Biochim Biophys Acta, 1999. 1410(2): p. 147–58.

4. Pitceathly, R.D. and R. McFarland, Mitochondrial myopathies in adults and children: management and therapy development. Curr Opin Neurol, 2014. 27(5): p. 576–82.

5. Schaefer, A.M., R.W. Taylor, D.M. Turnbull, and P.F. Chinnery, The epidemiology of mitochondrial disorders--past, present and future. Biochim Biophys Acta, 2004. 1659(2-3): p. 115–20.

6. Picard, M., J. Zhang, S. Hancock, O. Derbeneva, R. Golhar, P. Golik, S. O’Hearn, S. Levy, P. Potluri, M. Lvova, et al., Progressive increase in mtDNA 3243A>G heteroplasmy causes abrupt transcriptional reprogramming. Proc Natl Acad Sci U S A, 2014. 111(38): p. E4033–42.

7. Baughman, J.M., R. Nilsson, V.M. Gohil, D.H. Arlow, Z. Gauhar, and V.K. Mootha, A computational screen for regulators of oxidative phosphorylation implicates SLIRP in mitochondrial RNA homeostasis. PLoS Genet, 2009. 5(8): p. e1000590.

8. Aras, S., H. Arrabi, N. Purandare, M. Huttemann, J. Kamholz, S. Zuchner, and L.I. Grossman, Abl2 kinase phosphorylates Bi-organellar regulator MNRR1 in mitochondria, stimulating respiration. Biochim Biophys Acta Mol Cell Res, 2017. 1864(2): p. 440–448.

9. Aras, S., M. Bai, I. Lee, R. Springett, M. Huttemann, and L.I. Grossman, MNRR1 (formerly CHCHD2) is a bi-organellar regulator of mitochondrial metabolism. Mitochondrion, 2015. 20: p. 43–51.

10. Liu, Y., H.V. Clegg, P.L. Leslie, J. Di, L.A. Tollini, Y. He, T.H. Kim, A. Jin, L.M. Graves, J. Zheng, et al., CHCHD2 inhibits apoptosis by interacting with Bcl-x L to regulate Bax activation. Cell Death Differ, 2015. 22(6): p. 1035–46.

11. Aras, S., O. Pak, N. Sommer, R. Finley, Jr., M. Huttemann, N. Weissmann, and L.I. Grossman, Oxygen-dependent expression of cytochrome c oxidase subunit 4-2 gene expression is mediated by transcription factors RBPJ, CXXC5 and CHCHD2. Nucleic Acids Res, 2013. 41(4): p. 2255–66.

12. Aras, S., N. Purandare, S. Gladyck, M. Somayajulu-Nitu, K. Zhang, D.C. Wallace, and L.I. Grossman, Mitochondrial Nuclear Retrograde Regulator 1 (MNRR1) rescues the cellular phenotype of MELAS by inducing homeostatic mechanisms. Proc Natl Acad Sci U S A, 2020. 117(50): p. 32056–32065.

13. Purandare, N., N. Gomez-Lopez, M. Arenas-Hernandez, J. Galaz, R. Romero, Y. Xi, A.M. Fribley, L.I. Grossman, and S. Aras, The MNRR1 activator nitazoxanide abrogates lipopolysaccharide-induced preterm birth in mice. Placenta, 2023. 140: p. 66–71.

14. Stockis, A., X. Deroubaix, R. Lins, B. Jeanbaptiste, P. Calderon, and J.F. Rossignol, Pharmacokinetics of nitazoxanide after single oral dose administration in 6 healthy volunteers. Int J Clin Pharmacol Ther, 1996. 34(8): p. 349–51.

15. Koyano, F., K. Okatsu, H. Kosako, Y. Tamura, E. Go, M. Kimura, Y. Kimura, H. Tsuchiya, H. Yoshihara, T. Hirokawa, et al., Ubiquitin is phosphorylated by PINK1 to activate parkin. Nature, 2014. 510(7503): p. 162–6.

16. Purandare, N., Y. Kunji, Y. Xi, R. Romero, N. Gomez-Lopez, A. Fribley, L.I. Grossman, and S. Aras, Lipopolysaccharide induces placental mitochondrial dysfunction in murine and human systems by reducing MNRR1 levels via a TLR4-independent pathway. iScience, 2022. 25(11): p. 105342.

17. Walker, M.A., N. Slate, A. Alejos, S. Volpi, R.S. Iyengar, D. Sweetser, K.B. Sims, and J.E. Walter, Predisposition to infection and SIRS in mitochondrial disorders: 8 years’ experience in an academic center. J Allergy Clin Immunol Pract, 2014. 2(4): p. 465–468, 468 e1.

18. van Gisbergen, M.W., K. Offermans, A.M. Voets, N.G. Lieuwes, R. Biemans, R.F. Hoffmann, L.J. Dubois, and P. Lambin, Mitochondrial Dysfunction Inhibits Hypoxia-Induced HIF-1alpha Stabilization and Expression of Its Downstream Targets. Front Oncol, 2020. 10: p. 770.

19. Diaz-Trelles, R., M.C. Scimia, P. Bushway, D. Tran, A. Monosov, E. Monosov, K. Peterson, S. Rentschler, P. Cabrales, P. Ruiz-Lozano, et al., Notch-independent RBPJ controls angiogenesis in the adult heart. Nat Commun, 2016. 7: p. 12088.

20. Xie, C., X. Gao, D. Sun, Y. Zhang, K.W. Krausz, X. Qin, and F.J. Gonzalez, Metabolic Profiling of the Novel Hypoxia-Inducible Factor 2alpha Inhibitor PT2385 In Vivo and In Vitro. Drug Metab Dispos, 2018. 46(4): p. 336–345.

21. Fujita, N., D. Markova, D.G. Anderson, K. Chiba, Y. Toyama, I.M. Shapiro, and M.V. Risbud, Expression of prolyl hydroxylases (PHDs) is selectively controlled by HIF-1 and HIF-2 proteins in nucleus pulposus cells of the intervertebral disc: distinct roles of PHD2 and PHD3 proteins in controlling HIF-1alpha activity in hypoxia. J Biol Chem, 2012. 287(20): p. 16975–86.

22. Appelhoff, R.J., Y.M. Tian, R.R. Raval, H. Turley, A.L. Harris, C.W. Pugh, P.J. Ratcliffe, and J.M. Gleadle, Differential function of the prolyl hydroxylases PHD1, PHD2, and PHD3 in the regulation of hypoxia-inducible factor. J Biol Chem, 2004. 279(37): p. 38458–65.

23. Miikkulainen, P., H. Hogel, F. Seyednasrollah, K. Rantanen, L.L. Elo, and P.M. Jaakkola, Hypoxia-inducible factor (HIF)-prolyl hydroxylase 3 (PHD3) maintains high HIF2A mRNA levels in clear cell renal cell carcinoma. J Biol Chem, 2019. 294(10): p. 3760–3771.

24. Taniguchi, C.M., E.C. Finger, A.J. Krieg, C. Wu, A.N. Diep, E.L. LaGory, K. Wei, L.M. McGinnis, J. Yuan, C.J. Kuo, et al., Cross-talk between hypoxia and insulin signaling through Phd3 regulates hepatic glucose and lipid metabolism and ameliorates diabetes. Nat Med, 2013. 19(10): p. 1325–30.

25. Li, X.W., R.Z. He, Y. Li, and Z.F. Ruan, Tizoxanide mitigates inflammatory response in LPS-induced neuroinflammation in microglia via restraining p38/MAPK pathway. Eur Rev Med Pharmacol Sci, 2020. 24(11): p. 6446–6454.

26. Lokhande, A.S. and P.V. Devarajan, A review on possible mechanistic insights of Nitazoxanide for repurposing in COVID-19. Eur J Pharmacol, 2021. 891: p. 173748.

27. He, X., W. Hu, F. Meng, and X. Li, Design, Synthesis, and Ph*armacokinetic Evaluation of O-Carbamoyl Tizoxanide Prodrugs*. Med Chem, 2022. 18(1): p. 140–150.

28. Neary, M., U. Arshad, L. Tatham, H. Pertinez, H. Box, R.K.R. Rajoli, A. Valentijn, J. Sharp, S.P. Rannard, G.A. Biagini, et al., Quantitation of tizoxanide in multiple matrices to support cell culture, animal and human research. J Chromatogr B Analyt Technol Biomed Life Sci, 2023. 1228: p. 123823.

29. Guo, S., F. Li, B. Wang, Y. Zhao, X. Wang, H. Wei, K. Yu, and X. Hai, Analysis of tizoxanide, active metabolite of nitazoxanide, in rat brain tissue and plasma by UHPLC-MS/MS. Biomed Chromatogr, 2020. 34(2): p. e4716.

30. Birmingham, A., L.M. Selfors, T. Forster, D. Wrobel, C.J. Kennedy, E. Shanks, J. Santoyo-Lopez, D.J. Dunican, A. Long, D. Kelleher, et al., Statistical methods for analysis of high-throughput RNA interference screens. Nat Methods, 2009. 6(8): p. 569–75.

31. Purandare, N., M. Somayajulu, M. Huttemann, L.I. Grossman, and S. Aras, The cellular stress proteins CHCHD10 and MNRR1 (CHCHD2): Partners in mitochondrial and nuclear function and dysfunction. J Biol Chem, 2018. 293(17): p. 6517–6529.

32. Zhang, L., S. Peng, X. Dai, W. Gan, X. Nie, W. Wei, G. Hu, and J. Guo, Tumor suppressor SPOP ubiquitinates and degrades EglN2 to compromise growth of prostate cancer cells. Cancer Lett, 2017. 390: p. 11–20.

33. Wang, Y., X. Li, W. Liu, B. Li, D. Chen, F. Hu, L. Wang, X.M. Liu, R. Cui, and R. Liu, MicroRNA-1205, encoded on chromosome 8q24, targets EGLN3 to induce cell growth and contributes to risk of castration-resistant prostate cancer. Oncogene, 2019. 38(24): p. 4820–4834.

